# Early developmental morphology reflects independence from parents in social beetles

**DOI:** 10.1101/187740

**Authors:** Kyle M. Benowitz, Madeline E. Sparks, Elizabeth C. McKinney, Patricia J. Moore, Allen J. Moore

## Abstract

The variation in degree of offspring dependence in parents where parental care has evolved is striking, from feeding independence at birth to complete dependence on parents for all nutritional resources. This presents an evolutionary puzzle. Why lose the ability to feed as a contingency when parents may die or abandon broods? Comparisons of altricial and precocial vertebrates suggest that there may be life-history and developmental costs to early independence^1-3^. The generality of this beyond vertebrates is unclear, but we can extend the comparison as invertebrate species also vary in the level of independence in early life-history stages. For example, larvae of several burying beetle species (*Nicrophorus*), a genus in which parents regurgitate pre-digested food to begging larvae, have lost the ability to self-feed thus creating complete parental dependency for first instars^4^. Here, we ask whether variation in dependency amongst burying beetles is related to heterochrony in development of a more complex morphological structures. We show that the rate of development and allometry of mandibles of precocial larvae that can self-feed from birth are the same as those in altricial larvae that cannot survive without parenting. Instead, self-feeding is associated with shape variation in mandibles. In altricial species first instar larvae have smooth mandibles, whereas in precocial species mandibles are serrated. Later instars, which can self-feed in all species, have serrated mandibles. Serrations on teeth generally function to “grip and rip”^5^, whereas smooth blades function more to puncture^6^, and broods of altricial but not precocial *Nicrophorus* larvae show evidence for siblicide. We therefore suggest that altricial first-instar mandibles function more as weapons than feeding tools when released from self-feeding. This study presents a novel coevolution between developmental timing and parenting potentially mediated by sibling competition.

---

Why have juveniles of many organisms evolved absolute dependency on their parents? Investigations into the evolution of altricial (high dependence on parents at birth) and precocial (low dependence on parents at birth) species in a variety of taxa have revealed few overarching conclusions about the factors leading to these conditions^7^. However, several broad patterns have emerged. First, altricial birth is associated with faster life history strategies; altricial species are smaller and grow more quickly than precocial species in birds^1^, fish^2^, and mammals^3^. Second, altricial species display heterochronic development of specific features likely related to independence as compared to precocial species. In particular, eyes and external morphology (feathers/fur/cuticle) display delayed development in birds, mammals, and insects^1,3,8-10^. These trends suggest that when stable parenting reduces the need for offspring trait functionality, relaxed selection^11^ leads to delayed development of traits to conserve resources that can be allocated elsewhere, potentially related to rapid growth^1^. Here, we ask whether similar developmental associations are present in a unique insect genus that exhibits striking and discrete interspecific variation in the dependence of offspring on parents.

All species of beetles in the genus *Nicrophorus* provide extensive and elaborate parental care by directly regurgitating partially digested carrion into the mouths of their begging offspring^12^, and offspring of all species benefit from receiving parental regurgitations^13,14^. Parenting behaviour is remarkably similar across species, including function, to the extent that cross-fostering between species is readily accomplished without fitness effects on the recipient offspring^15^. However, there is discrete variation between species in the duration of the dependency of larvae on these parental regurgitations^16^; some larvae can survive from hatching without parents, while others will not survive past the first instar if parents are not present to feed^4,17^. There are no major differences in feeding ecology between obligate and facultative care species^4,12^, making this an ideal system for comparative studies. The phylogeny of *Nicrophorus* and its closest relatives suggests that larvae from obligate care species have lost the function of self-feeding during their early developmental stages, and that this process has occurred several times independently^17-19^. Dependence on parenting in *Nicrophorus* displays opposing life-history patterns from those observed in birds, fish, and mammals. Larger species, which develop more slowly as eggs and larvae^15^, consistently display obligate care across the *Nicrophorus* phylogeny whereas the opposite pattern is seen in vertebrates^1-3^. This raises the question of whether the developmental patterns associated with dependence may be different in burying beetles than in vertebrate parental systems. To ask this question, we investigated the timing of development in larval mouthparts in obligate and facultative parental feeding burying beetle species.

We examined the size of larval mandibles in one altricial (obligate care) species, *N. orbicollis*, and one precocial (facultative care) species, *N. vespilloides* (Fig. 1a, b), to determine if burying beetles show evidence for altered rates of development related to self-feeding. If *N. orbicollis* showed delayed mandible development, this would support a cost to developing the physical machinery related to feeding ability, providing a potential evolutionary explanation for parental dependence. We found no evidence for such a delay. Relative first instar mandible length was similar between the species (Fig. 1c). Furthermore, allometric relationships of mandible and head size between *N. orbicollis* and *N. vespilloides* did not differ in first (F_1,53_ = 0.16, p = 0.69), second (F_1,50_ = 0.03, p = 0.86), or third (F_1,35_ = 0.06, p = 0.81) instar larvae (Fig. 1c). Therefore, we find no evidence that *N. orbicollis* mouthparts develop at a slower rate than those of *N. vespilloides*.

**Figure 1.**
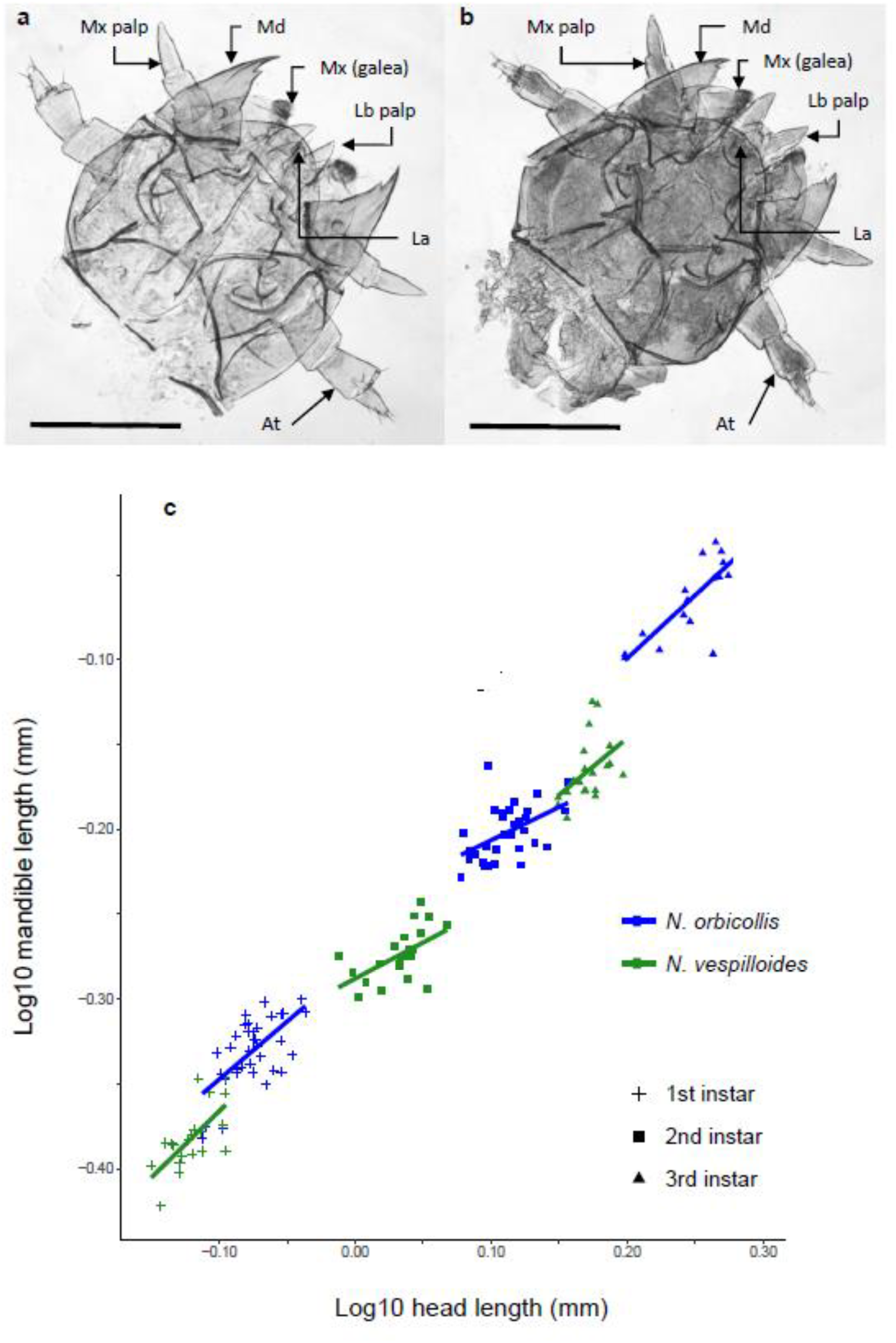
Mandible size in *N. orbicollis* and *N. vespilloides*. **ab,** Heads of first instar *N. orbicollis* (**a**) and *N. vespilloides* (**b**) larvae, with mouthparts and other appendages labelled. at, antenna; la, labrum, lb, labium; md, mandible; mx, maxilla. Scale bars, 500μm. **c,** Allometric relationships between mandible length and head length in first, second, and third instar *N. orbicollis* and *N. vespilloides*.

Although mandible size was not different, we found considerable variation in mandible shape between first instars of the altricial *N. orbicollis* and the precocial *N. vespilloides*. The interior edge of *N. vespilloides* first and second instar mandibles contained numerous jagged serrations (Fig. 2a, b), as seen in third instars of all Nicrophorine larvae^20^. In contrast, *N. orbicollis* first instar mandibles were completely smooth apart from a single major tooth on the interior edge (Fig. 2c). Furthermore, when *N. orbicollis* moult into the second instar and gain the ability to self-feed, their mandibles concurrently develop serrations like those seen in first and second instar *N. vespilloides* (Fig. 2d). In late first instar larvae, collected just before moulting, the serrated mandible can be seen developing underneath the cuticle of the smooth mandible (Fig. 2e). Serrations increase the ability to eat meat across a range of animal taxa, including dinosaurs, mammals, and insects^5^. Amongst insects, the presence of serrated structures is found on the mandibles of true bugs^21^, moths^22^, rove beetles^23^, and the ovipositors of *Drosophila suzukii*^24^, improving cutting ability. Here, serrated mandibles in the first instar likely allow *Nicrophorus* larvae to manipulate, tear and consume pieces of flesh from vertebrate carcasses upon which they feed and develop. Once serrations arise, they are retained in later instars, all of which self-feed and consume carrion that has not been processed by adults.

**Figure 2.**
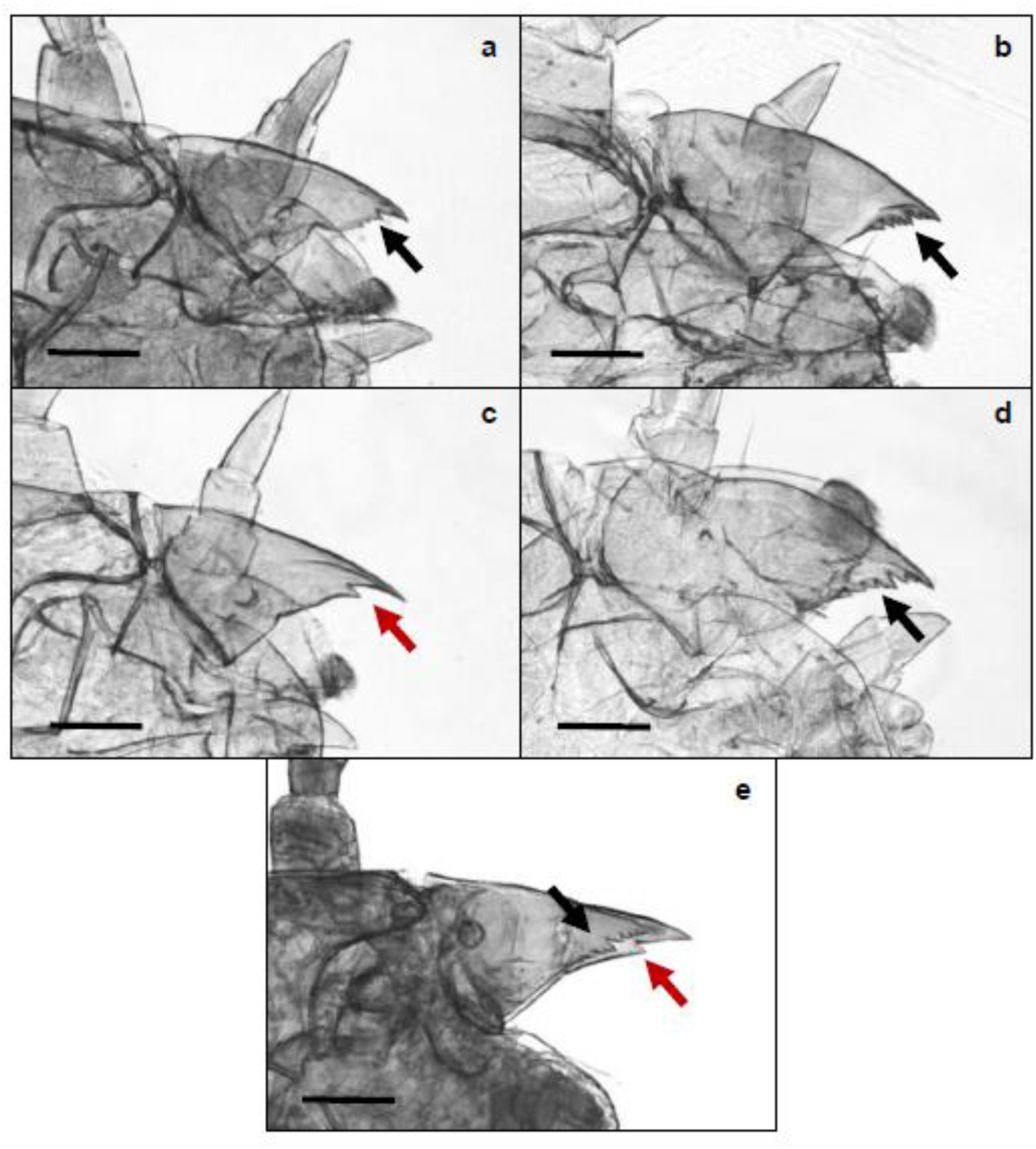
Mandible shape in *N. vespilloides* and *N. orbicollis*. **a,b** *N. vespilloides* first **(a)** and second **(b)** instar mandible showing the presence of serrations on the inner, cutting edge. **c,** *N. orbicollis* first instar mandible lacking serrations. **d,** *N. orbicollis* second instar mandible, with developed serrations above and below the initial large tooth. **e,** *N. orbicollis* late first instar mandible where developing second instar structure can be seen underneath the cuticle. Scale bars, 250μm. Arrows indicate the location of the inner mandible edge, red showing smooth, black showing serrated.

Most species of burying beetle facultatively self-feed, and few species where larvae are completely dependent are available; however, to further confirm the relationship between mandibular serrations and self-feeding ability we examined larvae of two more facultative *Nicrophorus* species and the other known obligate species. First instar larvae of precocial species *N. defodiens* and *N. tomentosus*^16^ were found to have mandibular serrations (Fig. 3a, b), consistent with the ability of these larvae to self-feed. However, in the altricial *N. sayi*^16^, both first and second instar larvae had smooth mandibles (Fig. 3c, d) before developing serrated mandibles in the third instar (Fig. 3e). The evolution of obligate care is thought to be phylogenetically independent in *N. orbicollis* and *N. sayi*^17,18^. Therefore, the loss of mandibular teeth appears to co-evolve with the loss of self-feeding ability in burying beetle larvae, and this morphology has evolved independently at least twice within the burying beetle lineage.

**Figure 3.**
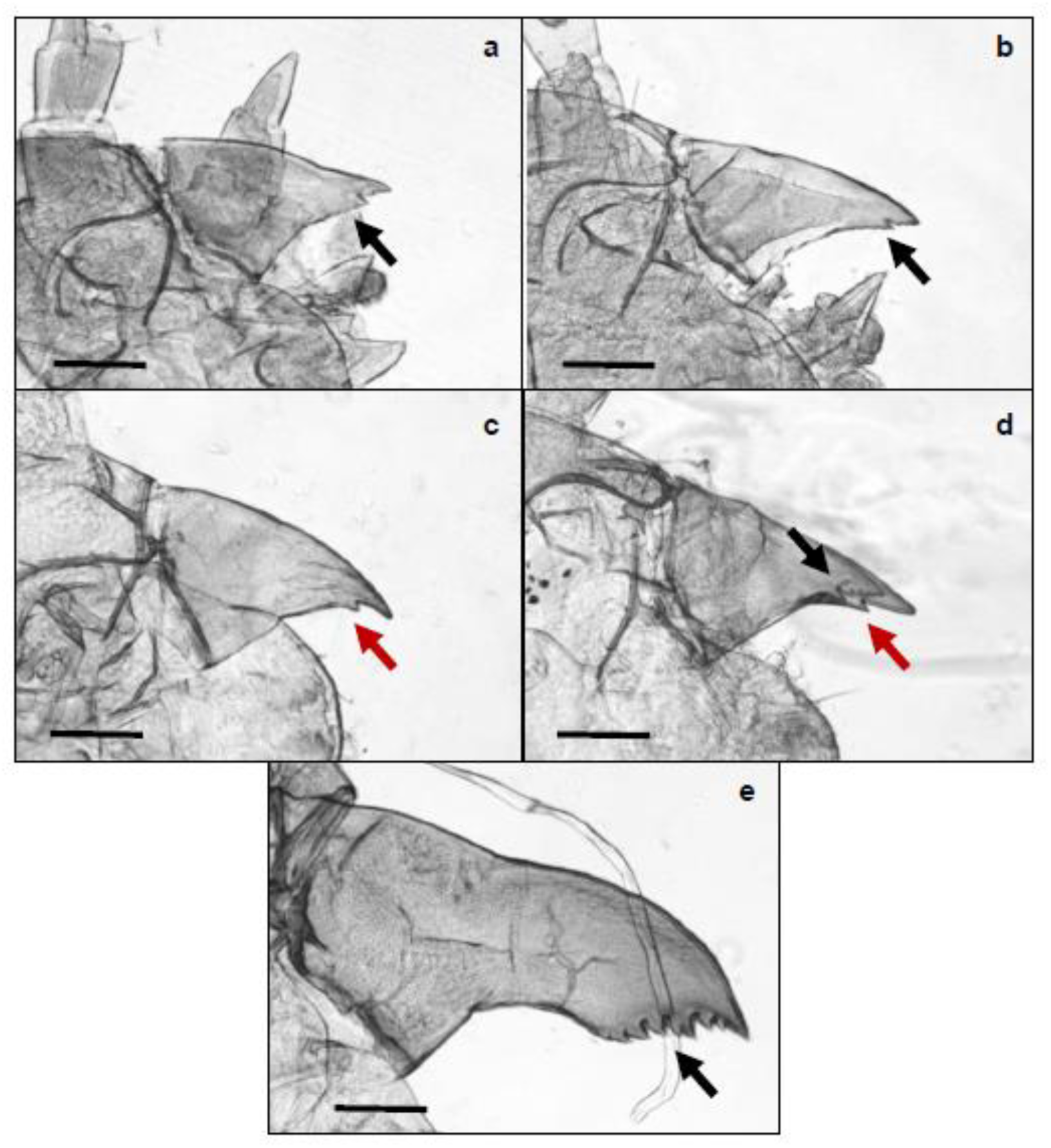
Mandible shapes of additional *Nicrophorus* species. **a,b** *N. defodiens* **(a)** and *N. tomentosus* **(b)** first instar mandibles, showing serrated inner edges as seen in *N. vespilloides.* **c,d,** First **(c)** and second **(d)** instar mandibles of *N. sayi*, lacking serrations, though serrated third instar mandible can be seen developing beneath the outer cuticle. **e** Third instar *N. sayi* mandibles, with pronounced serrations. Scale bars, 250μm. Arrows indicate the location of the inner mandible edge, red showing smooth, black showing serrated.

Our analyses suggest that the ability to self-feed does not incur a cost related to development, so what are the selective pressures leading to a morphology that facilitates self-feeding? Although there is evidence that *N. orbicollis* adults have relatively low mortality rates during reproduction and parenting^25^, the possibility of parental mortality or abandonment^26^ still exists. Self-feeding should be a strong strategy to deal with such contingencies. Accordingly, most species of burying beetles can self-feed at birth. Given this, we argue that there must be an additional functional benefit of having smooth mandibles. Specifically, mandibular variations could provide a competitive advantage to larvae in two ways: they could increase the efficiency of being fed by a parent, or increase the likelihood of being fed by a parent. In a previous experiment, we fostered *N. orbicollis* and *N. vespilloides* larvae with parents of both species^15^ and found that broods of *N. orbicollis* contained fewer larvae on average than broods of *N. vespilloides* (Extended Data Fig. 1), regardless of parent. This suggests that *N. orbicollis* undergoes a reduction in brood size in addition to parental culling that is not present in *N. vespilloides*, indicating the possibility of siblicide. Might this be related to the presence of smooth, sharpened mandibles? Larvae of some parasitoid wasps also use sharpened mandibles as weapons in siblicide^27^. Across animals, this is consistent with usage of smooth horns as stabbing weapons used in intraspecific combat in bovids^28^ as well as the use of smoothened teeth as piercing structures in sharks^6^. We therefore propose that *N. orbicollis* (along with other obligate care species) may be using smooth mandibles as weapons in sibling combat to reduce brood size and increase parental attention. However, the evolution of this behaviour may have necessitated the loss of a morphological trait, serrations, that allowed for feeding ability.

This suggests a novel roadmap for how offspring can evolve total dependence upon their parents. If parenting is relatively dependable, offspring may be released from constraint and able to evolve new functions using morphological structures previously necessary for feeding. In this way, an unexpected response to relaxed selection leads to coevolution between parenting behaviour and developmental timing of complex larval morphology.

## Methods

We collected *N. orbicollis* and *N. vespilloides* from Athens, GA and Cornwall, UK and maintained them at the University of Georgia as described previously^15^. We collected *N. defodiens* and *N. tomentosus* from Oneida County, WI^29^ in September 2016 and maintained them under identical conditions. We collected *N. sayi* from Oneida County and Vilas County, WI^29^ in May 2017 and maintained them at 15°C and a 16:8 light:dark cycle. We bred all beetles under these conditions by placing a male and a female in a plastic box (Pioneer Plastics, Dixon, KY, USA) and an 18-25g mouse carcass (RodentPro, Evansville, IN, USA). We collected first, second, and third instar larvae of *N. orbicollis*, *N. vespilloides*, and *N. sayi*, and first instar larvae of *N. defodiens* and *N. tomentosus*. We preserved collected larvae in 75% ethanol and stored them at 4°C until dissection.

To examine mandible size in *N. orbicollis* and *N. vespilloides*, we rinsed larvae twice in distilled water, removed the heads, and cleared the heads in 10% KOH. We then mounted each sample in KY Jelly (Reckitt Benckiser, Slough, UK), manually spread the mandibles away from the head capsule, and photographed them at 4x magnification using a Leica M80 stereomicroscope (Leica, Wetzlar, Germany). We measured mandible length from base to tip as well as vertical head capsule length (Extended Data Fig. 2) using Leica Application Suite morphometric software (LAS V4.1). We averaged left and right mandibles for analysis as there was no difference between them. We used type I ANOVA to test the effects of species, head length, and their interaction on mandible length in first (*N. v.* N=21; *N. o.* N=35), second (*N. v.* N=23; *N. o.* N=29), and third (*N. v.* N=20; *N. o.* N=18) instars. We report results for the interaction term (head size and species were highly significant for all instars), which informs whether the species display allometric differences in mandible size. We performed statistical analysis in R 3.4.0.

We washed and cleared larvae as described above to examine mandible shape across all five species. We then separated the mandibles and mounted them in glycerol. When required, we removed mouthparts other than the mandibles to ensure good visualization of the mandible structure. Photographs of the cleared heads were taken using a Leica DNIRE2 inverted microscope (Leica, Wetzlar, Germany) with a Hamamatsu model C4742-95 digital camera (Hamamatsu, Japan). To improve depth of field, a series of 4 to 7 images were stacked using Helicon Focus software (Kharkiv, Ukraine). Brightness and contrast of the overall images were adjusted using Adobe Photoshop CC (v. 2017.0.1) and, where needed, background debris was digitally removed from the image.

## Acknowledgements

We thank F. Ebot-Ojong and A. Amukamara for assistance in beetle rearing; J. McHugh for advice on preparing larval samples for microscopy; J. Johnson for taking high resolution images; and K. Ortman and S. Petersen for allowing us to collect beetles at Kemp Natural Resources Station and the Northern Highlands American Legion State Forest, respectively. K.M.B. was funded by a Kirby and Jan Alton Graduate Fellowship. NSF grant IOS-1354358 (to A.J.M.) funded this research.

### Author contributions

K.M.B. and A.J.M conceived and designed the experiments. K.M.B and E.C.M collected larval samples. K.M.B., M.E.S., and P.J.M. performed dissections and microscopy. M.E.S. performed measurements for allometric analysis. K.M.B. analysed allometric data. K.M.B, P.J.M., and A.J.M. wrote the manuscript with input from all authors.

### Competing financial interests

The authors declare no competing financial interests.

---

**Extended Data Figure 1.**
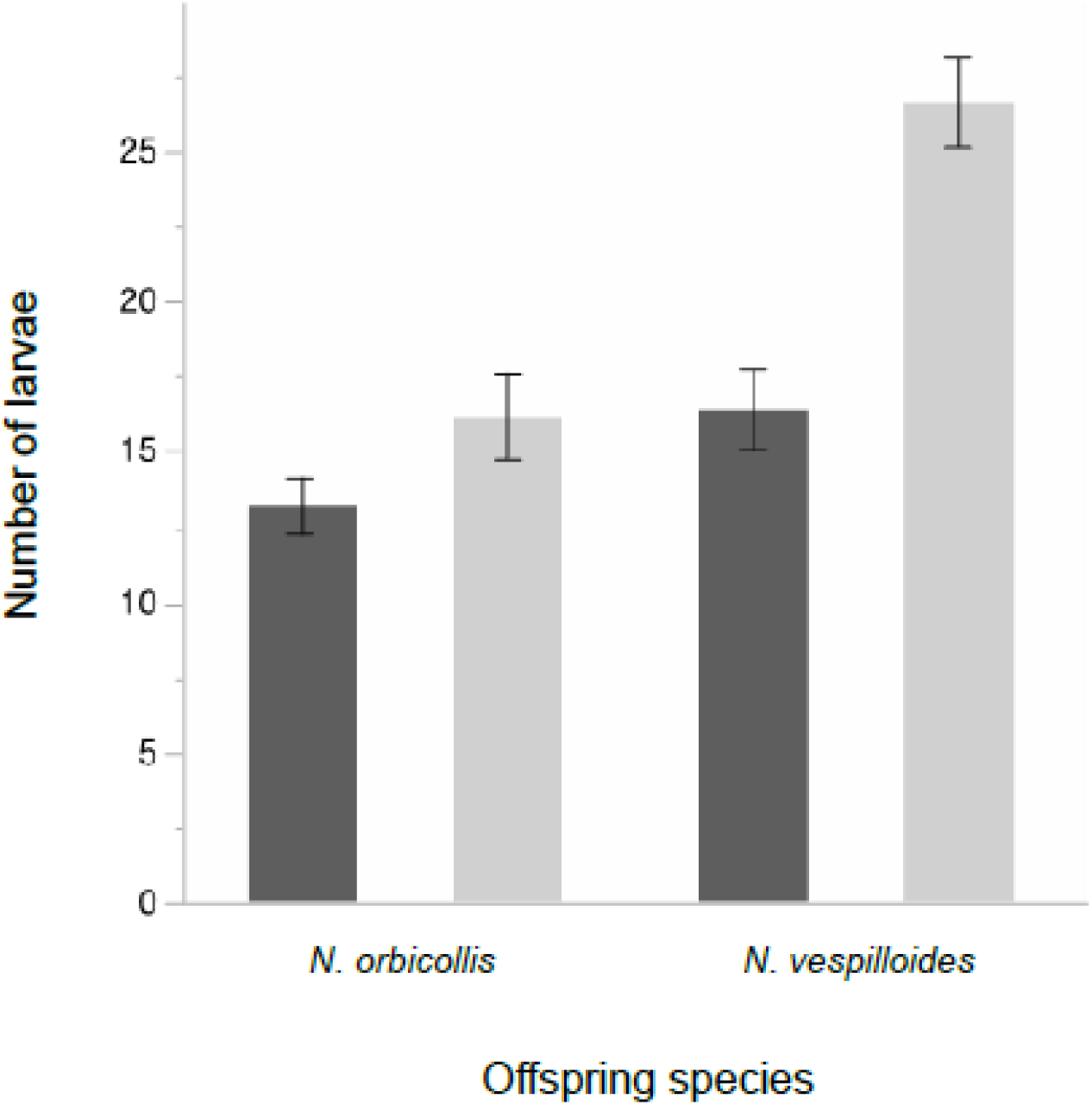
Number of larvae reared in cross-fostering between *N. orbicollis* and *N. vespilloides*. Dark bars indicate broods reared by *N. orbicollis* parents, light bars indicate broods reared by *N. vespilloides* parents. Unpublished data from Benowitz *et al.* 2015^15^.

**Extended Data Figure 2.**
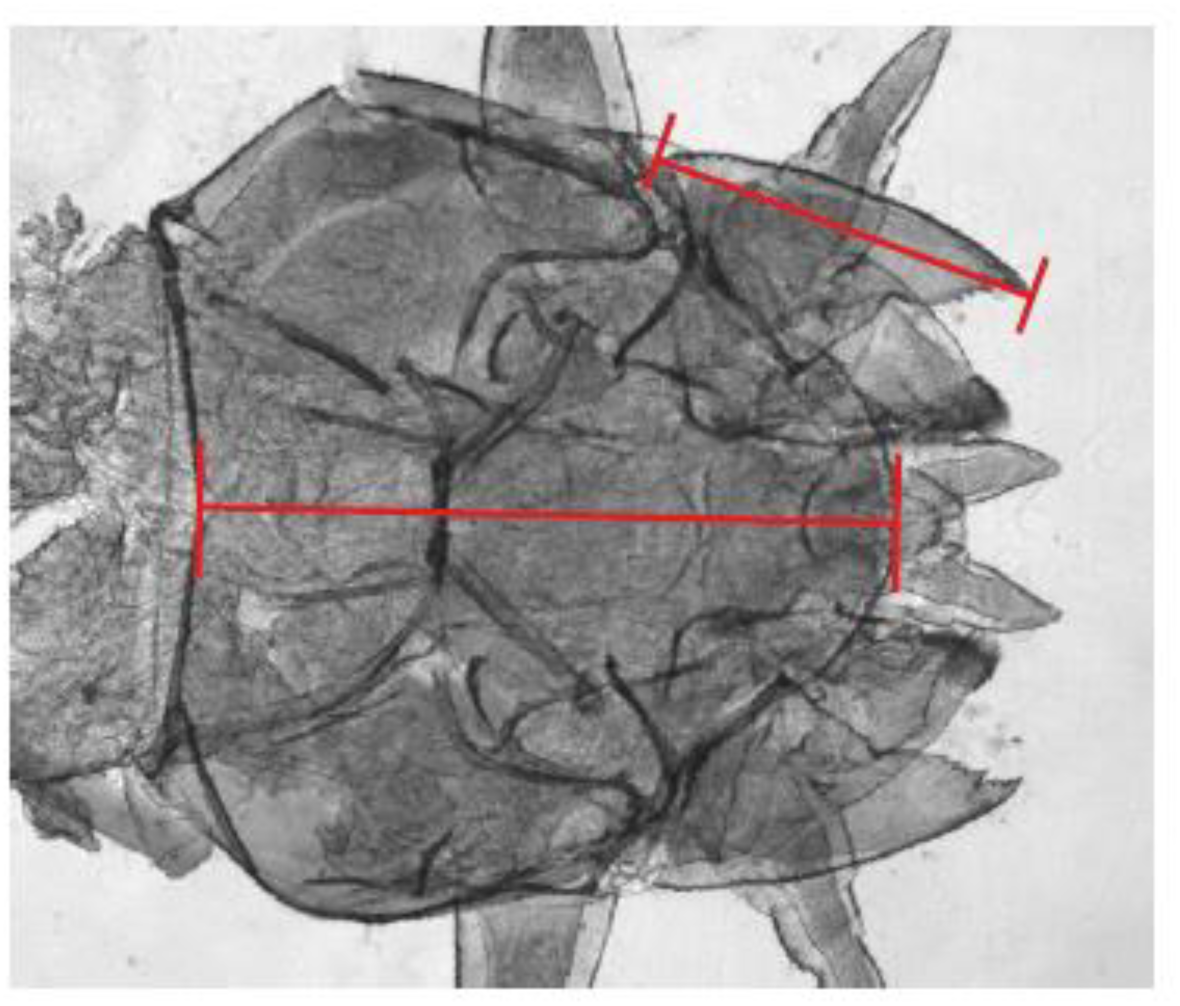
Measurement landmarks for mandible length and head capsule length. Mandibles were measured from the tip to the outer extreme of the base, whereas heads were measured down the centre line from the tip of the labrum to the posterior edge, as shown in red.

